# Allergic Airway Inflammation Protects Against SARS-CoV-2 and MERS-CoV Disease but Promotes Fatal SARS-CoV-1 Immunopathology

**DOI:** 10.64898/2026.09.02.748812

**Authors:** Jeremy Ardanuy, Grace Garrett, Carly Dillen, Meizi Liu, Louis Taylor, Achsah D. Keegan, Michael S. Diamond, Matthew B. Frieman

## Abstract

Epidemiological, clinical, and experimental data suggest that allergic asthma may have protective effects against severe Coronavirus Disease 2019 (COVID- 19) but are often conflicting between different models of infection and epidemiological analyses. Allergic asthma is characterized by type 2 (T2) inflammation, mucus hypersecretion, and airway remodeling. We used a house dust mite (HDM) respiratory exposure model to determine how preexisting allergic inflammation alters the pathogenesis of three highly pathogenic coronaviruses, SARS-CoV-1, SARS-CoV-2, and MERS-CoV. HDM treatment induced canonical features of allergic asthma, including goblet cell hyperplasia, mucin hypersecretion, eosinophil recruitment, M2 macrophage polarization, and induction of IL-13. Across multiple SARS-CoV-2 variants, HDM treated mice exhibited reduced viral titers, less weight loss, diminished lung immunopathology, and improved survival rates. Similar protection was observed in a mouse model of MERS-CoV, with reduced lethality, lower viral burden, and blunted cytokine storm signatures in HDM treated mice. In contrast, allergic airway inflammation exacerbated SARS-CoV-1 pathogenesis, as HDM-treated mice developed fulminant alveolar inflammation, severe weight loss, elevated pro-inflammatory cytokines and chemokines, and increased mortality despite having reduced virus titer. Thus, in mice, allergic airway inflammation exerts opposing virus specific effects on coronavirus pathogenesis. These results reconcile conflicting epidemiological and mechanistic studies across coronavirus epidemics and highlight the need to consider host allergic status as a key variable influencing disease outcome from emerging coronaviruses.

## Introduction

Asthma, an airway disorder affecting 200 to 300 million people globally [1], is characterized by chronic inflammation, airway obstruction and hyperresponsiveness, and mucus hypersecretion [2]. Asthma can be divided into multiple endotypes with the most common form being allergic asthma. In humans, allergic asthma is typically defined by a type 2 (T2) inflammatory response, dominated by elevated levels of interleukin (IL)-4, IL-5, IL-13, as well as alarmin cytokines IL-25, IL-33, and thymic stromal lymphopoietin (TSLP) [3, 4]. Allergic asthma is associated with attacks triggered by house dust mites (HDM), cockroaches, or smoke [5, 6]. Roughly 50% of asthmatic adults and 90% of asthmatic children are classified as allergic asthmatics, making it the most prevalent clinical form [7, 8]. The relationship between allergic airway inflammation and respiratory virus infections is complex, with many but not all viral agents causing asthma exacerbations [9]. Rhinovirus, influenza virus, and respiratory syncytial virus exacerbate lung inflammation and asthma related symptoms in those with pre-existing asthma [9]. The relationship is bidirectional, as pre-existing asthma can also influence the outcome of respiratory viral infections through altered antiviral immune responses including impaired interferon responses [10, 11].

In the context of Betacoronavirus infections, including Severe Acute Respiratory Syndrome coronavirus (SARS-CoV)-1, SARS-CoV-2 (the etiologic agent of Coronavirus Disease 2019 (COVID-19)), and Middle Eastern Respiratory Syndrome coronavirus (MERS-CoV) [12], several comorbidities have been associated with disease severity and mortality. Older age, diabetes, hypertension, and obesity are strong predictors of poor outcomes, including higher fatality rates and hospitalizations for all three highly pathogenic viruses [13–17].

The COVID-19 pandemic has enabled large scale epidemiological and mechanistic study of asthma in the context of SARS-CoV-2 infection and disease. Several meta-analyses and large cohort studies concluded that allergic asthma is not associated with more severe COVID-19 outcomes [18–20]; rather, the opposite appears to occur [21]. Asthmatic pediatric populations show a reduced rate of SARS-CoV-2 infection and disease severity [22]. In children with asthma that were infected with SARS-CoV or MERS-CoV, there was no reported increase in virus-induced symptoms [13, 14], although these studies had small cohorts due to a substantially lower prevalence of infection. COVID- 19 patients with non-allergic asthma that require inhaled corticosteroids had no elevated risk of severe COVID-19 but often present with potentially confounding comorbidities including chronic obstructive pulmonary disease that are difficult to untangle [23].

Mechanistically, allergic asthma alters host immune pathways known to be involved in SARS-CoV-1 and SARS-CoV-2 pathogenesis. Allergic asthma skews toward a T2 immune response characterized by an upregulation of IL-13. Addition of IL-13 has been shown to reduce expression of ACE2, the receptor for SARS-CoV-1 and SARS-CoV-2, on airway epithelial cells [21] but upregulate a cofactor for viral entry, the serine protease TMPRSS2 [21, 22]. The addition of IL-13 to airway liquid interface cultures (ALI) also led to a reduction in SARS-CoV-2 viral titer [23, 24], and this effect was abrogated when mucus was removed, indicating an important role for both IL-13 and mucus [25]. Nonetheless, these results appear to contrast with other studies in mice, which showed that IL-13 treatment before SARS-CoV-2 infection is associated with greater lung infection, worse disease, and higher lethality rates [26, 27]. The differences in outcomes between Th2 cytokine addition and allergic models, such as HDM administration, demonstrate that proper physiological context is needed to model the complex lung environment of allergic airway inflammation [28].

Here, in a mouse model of allergic asthma, we demonstrate substantial protection against SARS-CoV-2 and MERS-CoV infections, characterized by. reduced viral titers, weight loss, lung immunopathology, and mortality. In contrast, allergic asthma does not protect mice from SARS-CoV-1, instead resulting in increased weight loss and inflammatory chemokine and cytokine production despite having reduced virus titer. These results suggest that unique interactions between virus and host in the context of allergic airway inflammation can differentially alter coronavirus pathogenesis and disease outcome.

## Results

### A HDM model of allergic lung inflammation elicits T2 inflammation transcriptional response, mucus hypersecretion, and increased allergy associated cell types

To study the impacts of allergic asthma *in vivo*, we utilized an established mouse model of allergic asthma in which mice are treated with HDM over a period of two weeks resulting in markers of allergic asthma and remodeling of the pulmonary microenvironment [24]. Eight to 10-week-old BALB/c mice were intranasally treated with 50 μg of HDM or PBS on day 0 and every fifth day for a total of 4 treatments (**Fig. 1A**). Three days after the final HDM treatment, lungs were harvested for pre-infection characterization of allergic airway inflammation. RNA from lungs was purified and analyzed by RNA-sequencing and Ingenuity Pathway Analysis. Many allergic and airway hyperresponsiveness pathways were induced following the HDM treatment course correlating with known pathways activated by allergic asthma (**Fig. 1B**) and matching those found upregulated in asthma patients and mouse models [25]; these included mucus hypersecretion associated genes (*Muc5b*, *Muc5ac*, *Clca1*, *Slc26a4*, *Mmp8*), cytokines and chemokines associated with T2 inflammatory responses (*Cxcl5*, *Ccl8*, *Ccl17*, *Il33*), and genes associated with M2a macrophage polarization (*Chil4*, *Retnla*, *Retnlg*, *Chi3l1*) **(Fig. 1C)** [26]. A volcano plot of this data visualizes the differentially expressed genes in this analysis (**SI Appendix Fig. S1**).

**Figure 1.**
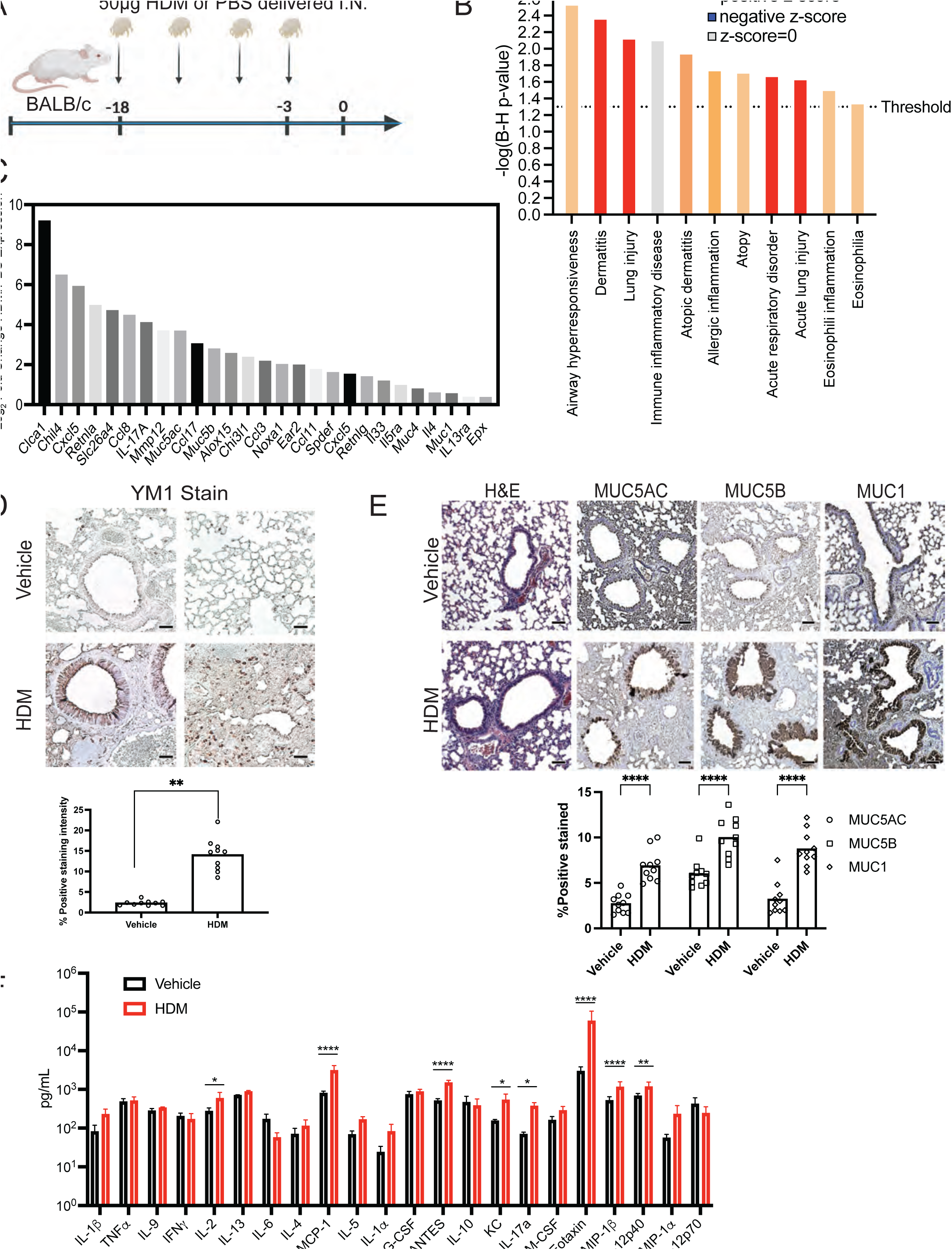
House dust mite administration induced allergic airway inflammation in a mouse model. (**A**) 8-week-old BALB/c mice were dosed four times with 50 µg of HDM at days -18, -13, -8, -3 prior to analysis. (**B**) Ingenuity pathway analysis of RNA-sequencing comparing HDM-dosed mouse lungs with control vehicle-dosed mouse lungs. (**C**) Top upregulated genes from RNA-sequencing comparison between HDM- and vehicle-dosed mouse lungs. (**D**) YM-1 IHC staining of mouse lungs for M2a macrophage population, quantification by % positive stained per field repeated 3 times per lung using ImageJ quantification. (**E**) Hematoxylin and eosin staining and IHC Mucin staining on mouse lungs, IHC quantification by % positive stained per field using ImageJ quantification. (**F**) Multiplex Luminex assay for chemokine and cytokine levels in mouse lungs. N=10 per group (data are pooled from two independent studies). Statistical analysis: Mann-Whitney U Test (**D**), Kruskal Wallis ANOVA (**E**) one way ANOVA with Tukey’s post-test used in (**F**) *P<0.05, **P<0.01, ***P<0.001, ****P<0.0001. Scale bars represent 200µM.

Macrophage polarization by IL-4 or IL-13 to an M2a phenotype is a hallmark of asthma pathogenesis [27, 28]. Immunohistochemical staining (IHC) for YM1, a marker of M2a macrophages was increased from 3% to 17% following induction of allergic conditions using HDM treatment as quantified by Image J analysis (**Fig. 1D**). Histological analysis of the lung revealed that HDM-treated mice have more peribronchiolar and alveolar space inflammation than vehicle-treated mice (**Fig. 1E**, *left*). Mucin hypersecretion leading to increased mucus in the lung is also a hallmark of allergic airway inflammation. Indeed, we observed increased levels of the mucins Muc5ac, Muc5B, and Muc1 in the HDM-treated mice from 3% to 11%, 4% to 14%, and 7% to 17% respectively (**Fig. 1E**).

We also assessed the effect of HDM treatment on cytokine and chemokine levels in the lung using a Luminex multiplex protein assay. This analysis revealed significant increases in IL-2, MCP-1, RANTES, KC, IL-17a, Eotaxin, MIP-1b, and IL-12 p40 levels (**Fig. 1F**). Higher chemokine levels were associated with the recruitment of multiple immune cell types including T cells, monocytes, neutrophils, and eosinophils as determined by flow cytometry and IHC analyses (**SI Appendix Fig. S1**). Together, these data demonstrate that the HDM treatment models allergic lung inflammation with mucus hypersecretion, increased allergic gene expressions, and accumulation of M2a macrophages and eosinophils.

### HDM treated mice are protected against severe COVID-19 following SARS-CoV-2 Beta variant infection

To determine whether pre-existing allergic airway inflammation affects SARS-CoV-2 infection and disease severity, 8–10-week-old BALB/c mice were dosed with HDM to induce allergic airway inflammation and inoculated 3 days after the final treatment with a low (10^2^ PFU) or high (10^5^ PFU) amount of SARS-CoV-2 B.1.351 (Beta variant) as described previously [29, 30]. Mice were weighed daily for 4 days to assess the effects on clinical disease state (**Fig 2A**). At the 10^2^ PFU challenge dose, the vehicle-treated mice had lost 5% of their weight by day 3, whereas the HDM-treated mice exhibited no significant weight loss. At the 10^5^ PFU dose, the vehicle-treated mice sustained ∼10% weight loss by day 3, In comparison, the HDM-treated mice infected with 10^5^ PFU of B.1.351 lost 8% of their starting weight but recovered by 3 dpi **(Fig 2A**). Lungs were harvested at 2 and 4 dpi for viral titer quantification, RNA isolation, and histological analysis. HDM-treated mice had approximately 30-fold reductions (*p* < .01) in lung viral titers compared to the vehicle-treated mice for both high and low dose challenges (**Fig 2B**).

**Figure 2.**
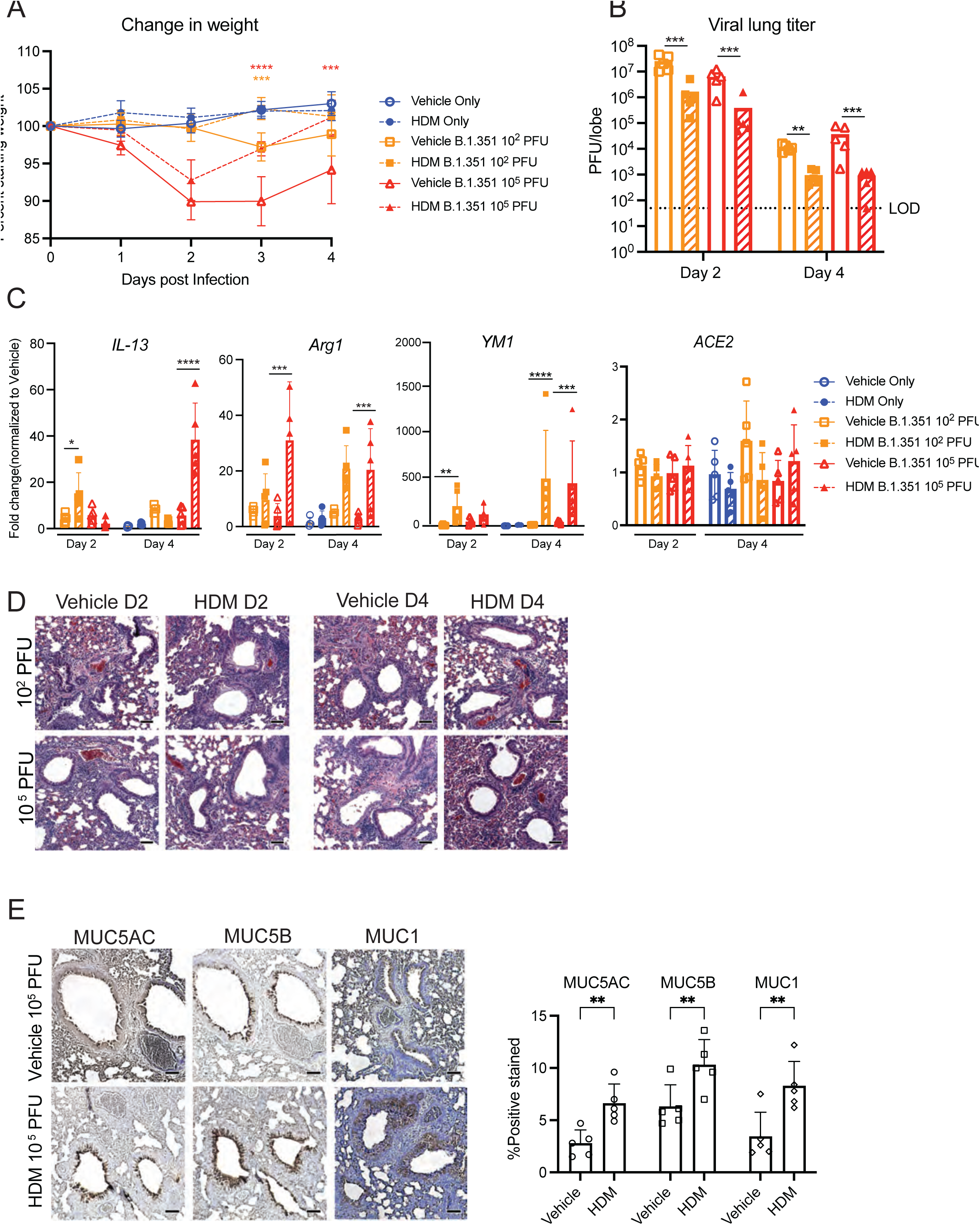
Allergic airway inflammation is protective in SARS-CoV-2 Beta variant challenge model. (**A**) 8-10-week-old BALB/c mice pretreated with four doses of HDM were inoculated with either high 10^5^ PFU (Red) or low 10^2^ PFU (Orange) of B.1.351 SARS-CoV-2 at day 0. Weights were recorded daily. (**B**) Lungs were harvested at 2 and 4 dpi, and infectious virus was quantified by plaque assay. (**C**) Lungs were analyzed for mRNA expression levels of *Il13*, *Arg1*, *Ym1*, and *Ace2* via qRT-PCR and normalized to uninfected vehicle treated group. (**D**) Lungs were fixed, embedded, sectioned, and stained by H&E to visualize lung inflammatory pathology as shown in representative images. (**E**) Lungs were fixed, embedded, sectioned and stained with IHC antibodies for indicated mucins. IHC quantification by % positive stained per field using ImageJ quantification. N=5 per group (data are representative of two independent studies). Statistical analysis: Two way- ANOVA test with Tukey’s multiple post-test (**A**) and Kruskal-Wallis ANOVA with Dunn’s post-test (**B, C, E**). *P<0.05, **P<0.01, ***P<0.001, ****P<0.0001. Scale bars represent 200µM.

We analyzed the host response to lung infection by evaluating changes in transcript levels of chemokines and cytokines by qPCR (**Fig. 2C**). Key cytokines of T2 inflammation were quantified. *Il13* levels in HDM-treated mice exhibited increased expression in both the low and high inocula infection groups, whereas the vehicle treated mice did not. Multiple markers of M2a macrophages, including *Chi3l3* and *Arg1*, were increased by 2 and 4 dpi in HDM-treated mice (**Fig. 2C**). We also evaluated ACE2 expression, the receptor for SARS-CoV-2 entry, since it is downregulated in epithelial cells formed during goblet cell hyperplasia [31] and could be responsible for the reduced virus levels in the lungs after infection of HDM-treated mice. However, *Ace2* expression was similar in HDM-treated and vehicle-treated mice (**Fig 2C**, *right*).

Lung histopathology was evaluated at 2 and 4 dpi (**Fig 2D**). In the HDM-treated groups, mice inoculated with the low dose of virus showed evidence of bronchiolar inflammation and inflammatory cell cuffing around the larger bronchi, as expected in an allergic asthma model [32], but lacked consolidation of alveolar spaces or alveolitis, this contrasts with the low dose vehicle group which had a lesser degree of bronchiolar inflammation and cuffing around the bronchi. In the high dose infection group, vehicle-treated mice displayed increased perivascular and peribronchiolar inflammation, including thickening around bronchi and denuding bronchiolitis by 2 dpi. With either viral dose, at 4 dpi, the HDM-treated and SARS-CoV-2-infected lungs had higher levels of infiltrating inflammatory cells, including eosinophils and monocytes, in the alveolar space. This indicates an overall increase of inflammation with SARS-CoV-2 infection in the context of HDM treatment; despite this, allergic inflammation specifically reduced the overall severity of viral disease.

One hypothesis as to how HDM-treated mice could limit infection from SARS-CoV-2 is through increased secretion of mucins that prevent SARS-CoV-2 virions from reaching the surface of target cells [33]. IHC analysis of mouse lungs at 4 dpi showed low levels of mucins MUC5AC, MUC5B, and MUC1 in the vehicle-treated infected mice, with staining concentrated around the epithelial surface of the airways (**Fig. 2E**). Both uninfected and infected HDM-treated mice had greater mucin staining in the airways, which might impact SARS-CoV-2 infection and spread. The mucin staining levels for MUC5AC, MUC5B, and MUC1 increased from 3 to 7%, 6 to 10%, and 3 to 8%, respectively

Collectively, these experiments demonstrate that HDM-treatment in mice leads to T2 immune skewing, like that seen in asthmatic humans with allergic asthma, and that this immune environment protects the lungs against severe disease caused by SARS-CoV-2.

### HDM treated mice are protected against infection by an Omicron variant and a mouse adapted SARS-CoV-2

To determine if the protective phenotype was consistent across multiple SARS-CoV-2 strains, we evaluated the pathogenesis of the SARS-CoV-2 Omicron variant BA.5 in the HDM allergic airway inflammation mouse model. BA.5 emerged in 2022 and caused increased transmission compared to the previous SARS-CoV-2 strains. Eight to 10-week-old BALB/c mice were treated with HDM or vehicle as described above before intranasal challenge with 10^5^ PFU of BA.5. After infection, vehicle-treated mice sustained ∼5% weight loss by 2 dpi and recovered to their starting weight by 4 dpi, whereas HDM-treated mice infected with BA.5 showed no appreciable weight loss, and instead, had weight gain at 2 and 3 dpi (**Fig. 3A**). Lungs from these mice were also evaluated for virus titer at 4 dpi. Compared to B.1.351, BA.5 and other Omicron variants replicate less efficiently in mice [34, 35], as evidenced by lower lung viral titers (∼10^3^ PFU) at 4 dpi in vehicle-treated mice. Lungs from BA.5 infected HDM-treated mice had a 10-fold (*p* < .01) reduction in viral titer compared to vehicle-infected mice (**Fig. 3B**) demonstrating a consistent phenotype of reduced SARS-CoV-2 infection titer in the lungs of HDM-treated mice. Histological evaluation revealed that HDM-treated and BA.5-infected mice had increased perivascular inflammation and thickening of the small airways compared to vehicle-treated BA.5 infected mice (**Fig. 3C**). Even after infection with a less pathogenic variant, a greater inflammatory response is present in the HDM-treated mice, while still having decreased weight loss and virus replication. Also as seen with B.1.351 infection, the vehicle-treated mice had lower levels of mucins after BA.5 infection (3%), than HDM-treated mice (18%) and HDM treated BA.5-infected mice (7%) (**Fig. 3D** and **SI Appendix, Fig. S2**).

**Figure 3.**
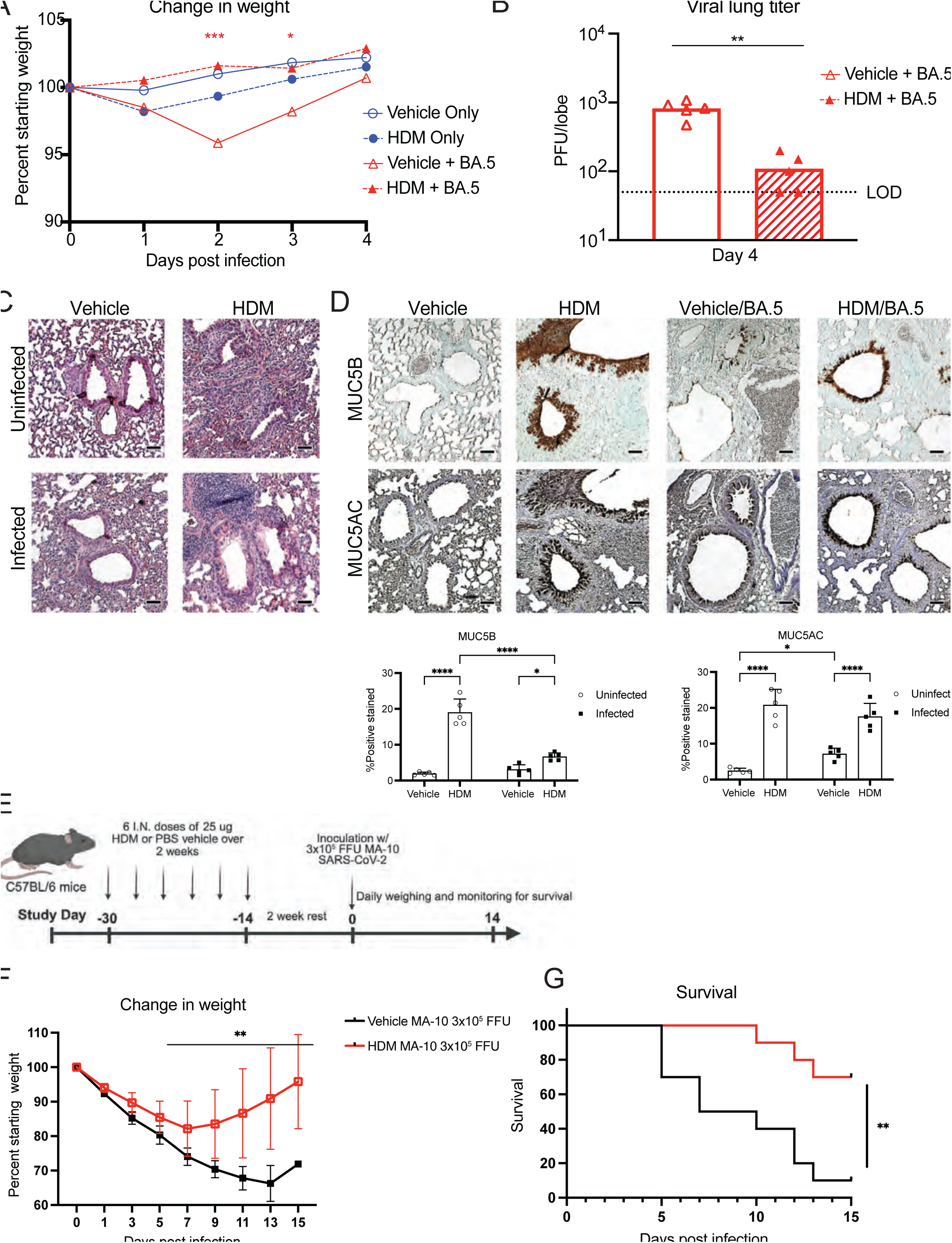
HDM-induced airway inflammation abrogates SARS-CoV-2 Omicron infection and reduces lethality of mouse adapted SARS-CoV-2. (**A**) 8-10-week-old BALB/c mice were dosed four times with HDM and then challenged with SARS-CoV-2 BA.5, and weight was recorded daily. (**B**) Lung viral titer was quantified at 4 dpi. (**C**) H&E staining on lung at 4 dpi to visualize lung inflammatory pathology. (**D**) IHC staining on lungs for Muc5B and Muc5AC at 4 dpi. IHC quantification by % positive stained per field using ImageJ quantification. (**E**) 20-week-old C57BL/6 mice were dosed with 6 x 25 µg of HDM over 2 weeks and then challenged 2 weeks later with 3 x 10^5^ focus-forming units (FFU) of MA-10. (**F**) Mice were weighed daily and recorded. (**G**) Survival and mortality were monitored daily. N=10 per group in MA-10 figures and N=5 per group in Omicron studies (data are representative of two independent studies). Statistical analysis: (**A**, **D, F**) Kruskal-Wallis ANOVA with Dunn’s post-test. (**B**) Mann-Whitney test. (**G**) Log-rank Mantel-Cox test. *P<0.05, **P<0.01, ***P<0.001, ****P<0.0001. Scale bars represent 200µM.

We next evaluated the effect of HDM treatment using a severe model of SARS-CoV-2, a mouse adapted strain (MA-10), and a different mouse genetic background. This strain of mice required a modified HDM dosing scheme [32] to determine if similar protection against lethal disease could be observed. Using 20-week-old C57BL/6 mice, six doses of 25 μg of HDM or PBS vehicle were administered intranasally over the course of two weeks to induce allergic airway inflammation. The mice were then rested for two weeks prior to intranasal challenge with 3 x 10^5^ FFU of MA-10 (**Fig. 3E**). Vehicle-treated, infected mice lost over 30% of their weight by 13 dpi and failed to rebound, whereas the peak weight loss of MA-10 infected and HDM-treated mice was 18% at 7 dpi before rebounding toward starting weights (**Fig. 3F**). This improvement in weight was accompanied by higher rates of survival following MA-10 infection: 70% of HDM-treated mice versus 10% of vehicle-treated mice (**Fig. 3G**).

### Allergic airway inflammation protects mice from lethal MERS-CoV infection

Unlike SARS-CoV-2, which efficiently infects epithelial cells throughout both the upper and lower respiratory tract, MERS-CoV exhibits a predominantly lower respiratory tract tropism due to the restricted expression of its cellular receptor, DPP4, on distal airway and alveolar epithelial cells [36]. To assess whether the reduction in pathogenesis in HDM-treated mice was broadly applicable to other coronaviruses, we tested for effects in a model of MERS-CoV pathogenesis using human DPP4 knock-in C57BL/6 mice and a mouse adapted MERS-CoV (MERS-MA) [37]. DPP4^KI/KI^ mice were treated with four doses of HDM or PBS over two weeks and then inoculated intranasally with 5 x 10^3^ PFU of MERS-MA. Consistent with previous data [29], vehicle-treated mice sustained greater than 20% weight loss starting at 2 dpi that worsened through 5 dpi. Conversely, HDM-treated mice infected with MERS-MA had minimal weight loss, losing only 4% starting weight by 3 dpi before fully recovering (**Fig. 4A**). Lethality was observed in the vehicle-treated groups as early as 3 dpi, with 60% mortality by 7 dpi, whereas no deaths occurred in the HDM-treated and MERS-MA infected mice (**Fig. 4B**).

**Figure 4.**
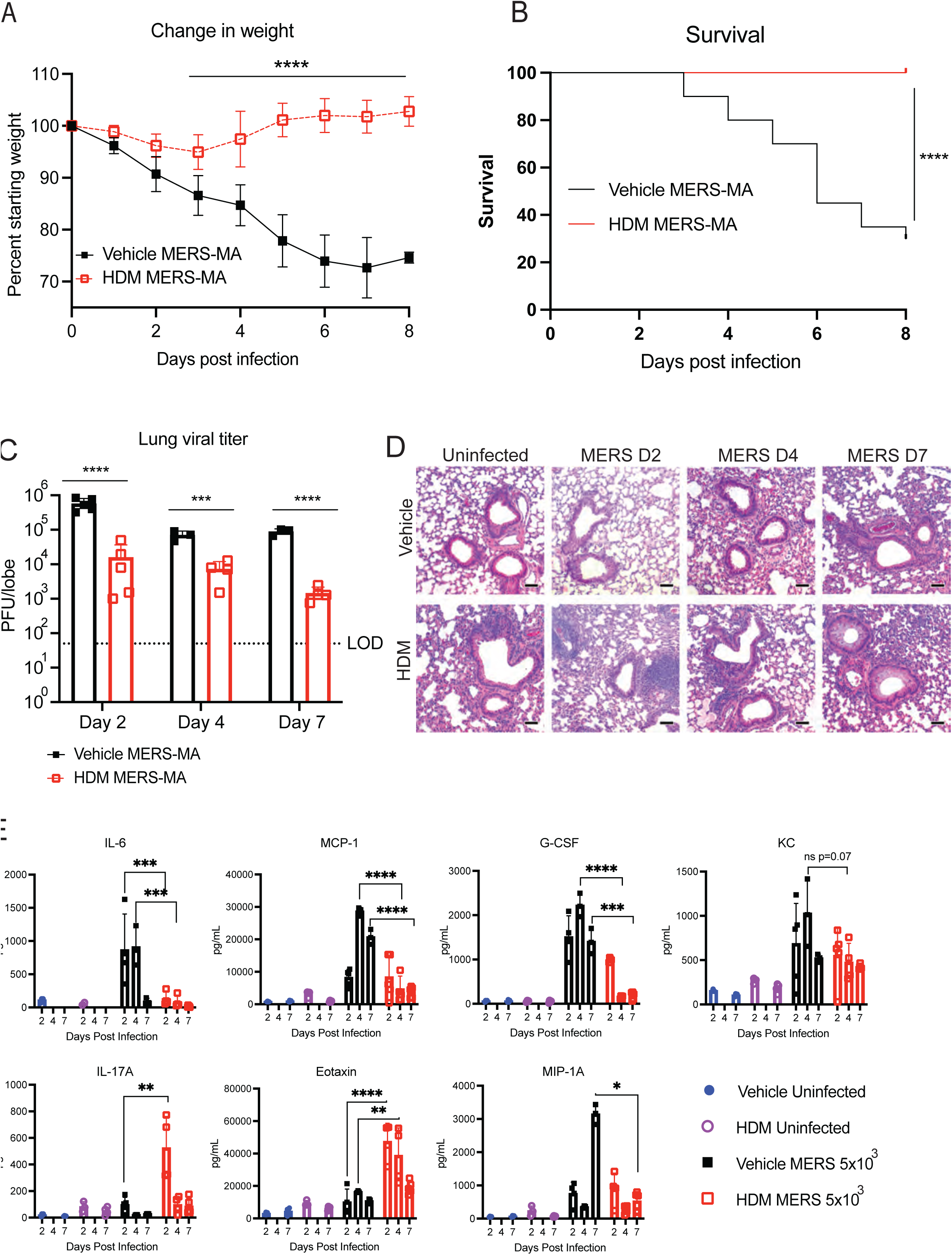
MERS-CoV disease and lethality is prevented by allergic airway inflammation. (**A**) 8-10-week-old humanized DPP4 KI/KI mice were treated with four 50-µg doses of HDM or vehicle and then challenged with MERS-MA three days later. Weights were measured daily. (**B**) Survival and mortality were recorded daily. (**C**) Lung viral titers were quantified by plaque assay at 2, 4, and 7 dpi. (**D**) Lungs were fixed, embedded, sectioned, and stained for H&E to visualize lung inflammatory pathology. (**E**) Luminex protein assay for cytokines and chemokines was from lung homogenates at 4 dpi. N=5-10 per group (data are representative of three independent studies). Statistical analysis: (**A**) Kruskal-Wallis ANOVA with Dunns post-test. (**B**) Log-rank Mantel-Cox test. (**C**) Mann-Whitney test. (**E**) Two-way ANOVA with Tukey’s post-test. *P<0.05, **P<0.01, ***P<0.001, ****P<0.0001. Scale bars represent 200µM.

Mice were euthanized at 2, 4, and 7 dpi, and lungs were collected for viral titer. In HDM-treated mice, 10 to 100-fold reductions (*p* < .01) in MERS-CoV viral titer were observed at all three timepoints compared to vehicle-treated mice (**Fig 4C**). Histological analysis showed that vehicle-treated, MERS-MA infected mice had little airway inflammation at 2 dpi, but by 4 and 7 dpi bronchiolar thickening and alveolitis were present, with high levels of myeloid cell recruitment. In the HDM-treated and MERS-MA infected mice, neutrophil and eosinophil recruitment was observed by 2 dpi, and this correlated with less interstitial inflammation at 4 and 7 dpi (**Fig. 4D**).

Lung homogenates from these time points were also evaluated for cytokine and chemokine levels (**Fig. 4E**). Key innate immune cell recruitment and tissue remodeling markers including MCP-1, G-CSF, and MIP-1a were all higher in vehicle-treated, infected mice than HDM-treated, infected mice. Additionally, IL- 17a and eotaxin were higher in HDM-treated and infected mice than vehicle-treated and infected mice, suggesting that T helper cells and eosinophils could be associated with protection in MERS-MA models of disease.

To determine whether hDPP4 gene dosage contributed to the protective phenotype, we repeated HDM treatment and MERS-MA infection in hDPP4^KI/+^ heterozygous mice. HDM-treated heterozygous mice exhibited similar protection from weight loss and lethality as hDPP4^KI/KI^ homozygous mice **(SI Appendix, Fig. S3**). Furthermore, lung DPP4 expression was unchanged following HDM treatment (**SI Appendix, Fig. S3**), indicating that altered receptor expression was not responsible for the observed protection.

Taken together, these data show that HDM treatment markedly reduces MERS-CoV pathogenesis, resulting in decreased weight loss, lethality, lung viral titers, and airway inflammation. These findings parallel those observed following SARS-CoV-2 infection, despite the viruses differing in receptor usage (DPP4 versus ACE2) and respiratory tract tropism.

### Allergic airway inflammation worsens SARS-CoV-1 disease and lethality in mice

There is limited data from the SARS-CoV-1 outbreak in 2002-2003 analyzing the relationship between allergic asthma and disease severity. As SARS-CoV- 1 uses the same receptor, ACE2, and has a similar cell tropism to SARS-CoV- 2, we hypothesized that HDM treatment might protect against SARS-CoV-1 infection. We implemented the same four dose HDM treatment over two weeks in BALB/c mice followed by infection with 10^5^ PFU of mouse adapted SARS-CoV-1 (MA-15). Mice were weighed daily and monitored for survival through 8 dpi. A subset of mice was euthanized at 4 dpi, and lungs were harvested for plaque assay, histopathology, and cytokine/chemokine quantification.

Consistent with previous experiments [29], MA15 infection of vehicle-treated mice lost ∼10% of their body weight by 4 dpi and rebounded to their starting weight by 8 dpi. In comparison, HDM-treated mice lost more weight, ∼30% of their starting weight by 5 dpi (**Fig. 5A**). Vehicle-treated, SARS-CoV-1 infected all survived infection, whereas 100% of HDM-treated, infected mice died by 6 dpi (**Fig. 5B**). Notwithstanding this result, HDM-treated mice had ∼10-fold lower (*p* < .01) lung viral titers than vehicle-treated mice (**Fig. 5C**).

**Figure 5.**
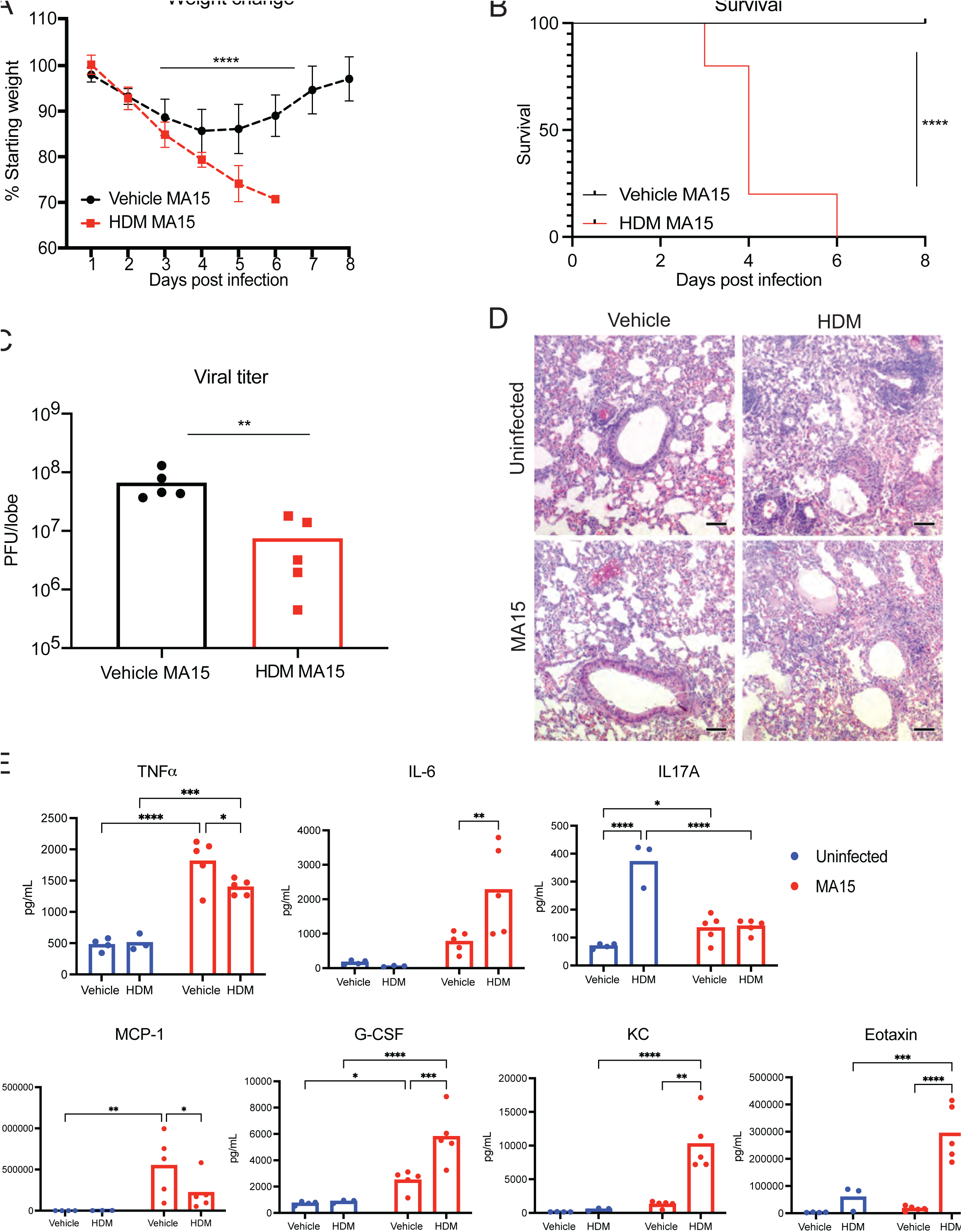
SARS-CoV-1 disease and lethality is exacerbated by allergic airway inflammation. (**A**) 8-10-week-old BALB/c mice were dosed with 4 x 50 µg of HDM and then challenged with 10^5^ PFU of SARS-CoV-1 MA-15. Weights were measured daily. (**B**) Survival and mortality were recorded daily. (**C**) Lung viral titer was quantified by plaque assay at 4 dpi. (**D**) Lungs were fixed, embedded, sectioned, and stained with H&E to visualize pathology. (**E**) Luminex protein assay for cytokines and chemokines was from lung homogenates at 4 dpi. N=10 per group (data are representative of two independent studies). Statistical analysis: (**A**) and (**E**) Kruskal-Wallis ANOVA with Dunns post-test. (**B**) Log-rank Mantel Cox test (**C**) Mann-Whitney test. *P<0.05, **P<0.01, ***P<0.001, ****P<0.0001. Scale bars represent 200µM.

Because of the apparent disparity between viral burden and disease outcome in HDM- and vehicle treated SARS-CoV-1 infected mice, we investigated effects on lung pathology and inflammation. Lungs sections were stained with H&E at 4 dpi. In the uninfected, HDM treated groups, an increased accumulation of monocytes and neutrophils was observed compared to vehicle treated mice. Following SARS-CoV-1 infection, HDM-treated mice had higher levels of alveolar inflammation and consolidation than vehicle-treated mice, consistent with their more severe disease (**Fig. 5D**).

Lung cytokine/chemokine levels were measured at 4 dpi (**Fig. 5E**). The increased disease severity observed in infected HDM-treated mice was associated with elevated IL-6, G-CSF, KC (CXCL1), and eotaxin, suggesting enhanced acute inflammatory and granulocyte-associated responses. In contrast, TNF-α and IL-17A were unchanged, and MCP-1 (CCL2), a key chemokine for inflammatory monocyte recruitment, was reduced. These findings indicate that HDM exposure does not broadly amplify all inflammatory pathways but instead selectively promotes an IL-6- and neutrophil-associated inflammatory program that might exacerbate SARS-CoV-induced lung injury Collectively, these data indicate that HDM treatment of mice results in worsened SARS-CoV-1 disease and outcomes in mice. The results of SARS-CoV-1 infection contrast with the protective effects that allergic airway inflammation has during MERS-CoV and SARS-CoV-2 infection, suggesting unique pathogenesis pathways for each virus in this context.

### SARS-CoV-1 infection in HDM-treated mice is characterized by a cytokine storm signature

To further probe the basis for the SARS-CoV-1 disease phenotype in HDM-treated mice, we implemented the same four dose HDM treatment over two weeks in BALB/c mice followed by infection with a lower dose (10^2^ PFU or 10^3^ PFU) of MA-15. In the 10^3^ PFU group, HDM-treated mice lost ∼15% of their body weight by 4 dpi whereas vehicle-treated mice lost 10%. In the 10^2^ PFU group, both HDM-treated and vehicle-treated mice only lost around ∼5% bodyweight and began to recover by 4 dpi (**Fig. 6A**). Cohorts of mice were euthanized at 2 dpi, and their lungs were titrated for infectious virus. The 10^3^ PFU group of HDM-treated mice had ∼3-fold reduction (*p* < .01) in viral titer compared to the 10^3^ PFU infected, vehicle-treated mice, whereas the 10^2^ PFU groups showed no statistically significant differences (**Fig. 6B**). At the 10^3^ PFU 10 dose of MA-15, vehicle treated mice all survived though 4 dpi, whereas 80% of the HDM-treated mice succumbed. All 10^2^ PFU infected mice, regardless of HDM treatment, survived to 4 dpi (**Fig. 6C**). Lungs were stained with H&E at 4 dpi for histological analysis. Increased recruitment of monocytes and neutrophils was observed in the HDM-treated uninfected groups. After infection, the 10^3^ PFU group of HDM-treated mice had higher levels of alveolar inflammation and consolidation as compared to the similarly infected, vehicle-treated group (**Fig. 6D**).

**Figure 6.**
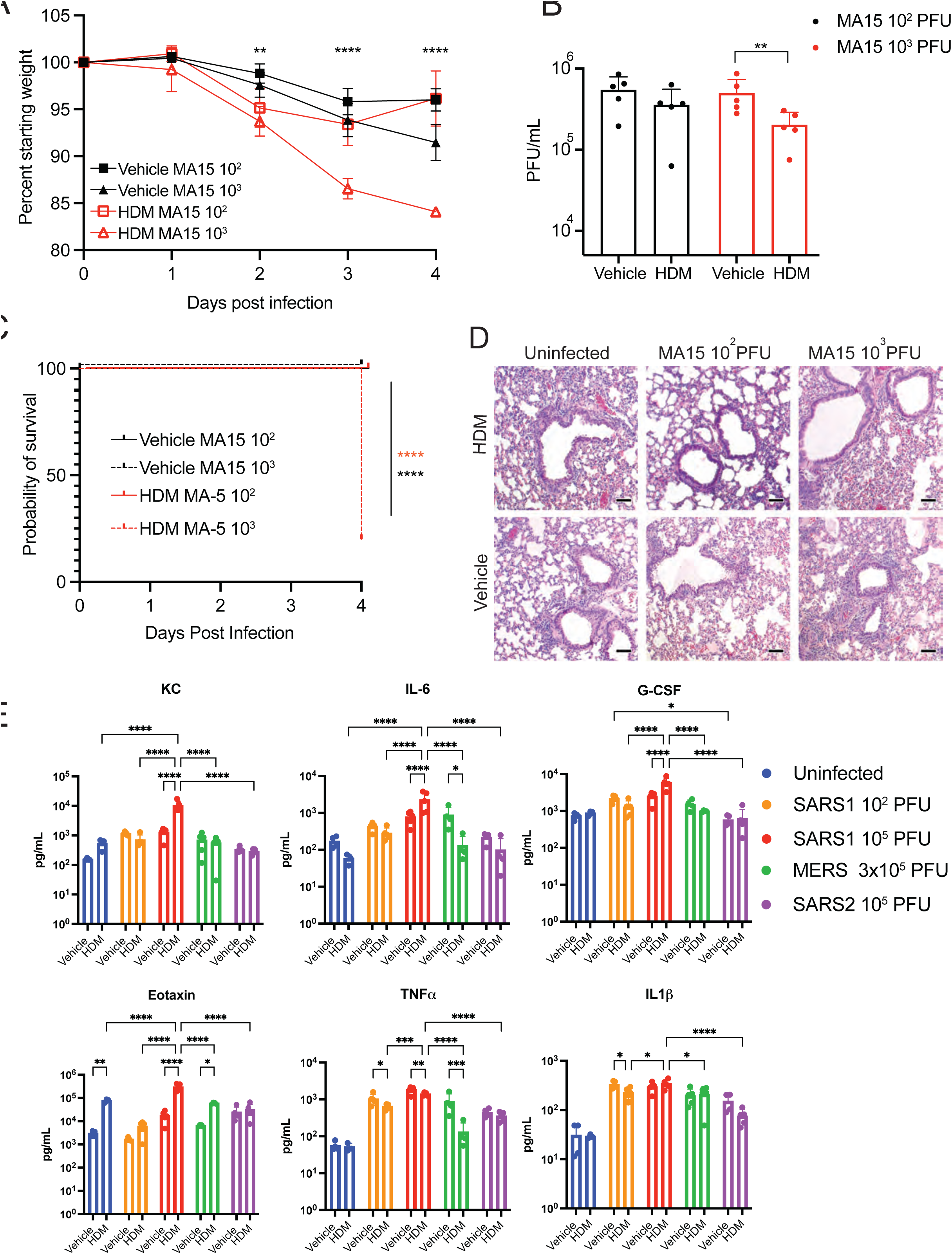
Allergic airway inflammation worsens SARS-CoV-1 disease and is characterized by increased proinflammatory cytokines. (**A**) 8-10-week- old BALB/c mice were dosed with 4 x 50 µg HDM and then challenged with SARS-CoV-1 10^2^ or 10^3^ MA-15. Weights were recorded daily. (**B**) Lung viral titers were quantified by plaque assay at 4 dpi. (**C**) Survival and mortality were recorded daily. (**D**) At 4 dpi, lungs were fixed, embedded, sectioned, and stained for H&E to visualize inflammatory pathology. (**E**) Luminex protein assay for cytokines and chemokines was from lung homogenates at 4 dpi. N=10 per group (data are pooled from two independent studies). Statistical analysis: (**A**) and (**E**) Kruskal-Wallis ANOVA with Dunn’s post-test. (**B**) Mann-Whitney test. (**C**) Log-rank Mantel Cox test. *P<0.05, **P<0.01, ***P<0.001, ****P<0.0001. Scale bars represent 200µM.

Mouse lungs at 4 dpi from the 10^2^ PFU groups were analyzed for chemokine and cytokine protein levels. These were compared with samples from previous experiments SARS-CoV-1 10^5^ MA-15 groups, MERS-CoV groups, and SARS-CoV-2 MA-10 groups all from 4 dpi for comparison (**Fig. 6E**). Notably, KC, IL-6, G-CSF, Eotaxin, TNF-α, and IL-1β were all increased in SARS-CoV-1 infected mice, with the HDM groups having higher levels of KC, IL-6, G-CSF, and Eotaxin. The proinflammatory cytokine and chemokine profile in the SARS-CoV- 1 infected, HDM-treated lungs is one associated with elevated neutrophils [38], which could promote disease. However, depletion of neutrophils using an anti- Ly6G antibody prior to infection did not protect mice from lethal SARS-CoV-1 challenge (**SI Appendix, Fig. S4**).

In summary, HDM pretreatment of mice skews immune responses towards a T2 mediated response that differentially affects the severity of disease caused by infection of several betacoronaviruses.

## Discussion

Understanding how pre-existing conditions, such as asthma and allergic airway inflammation, impacts the host responses to highly pathogenic coronaviruses has important implications for pandemic preparedness, resolving mechanisms of disease, treatment of distinct populations, and risk stratification of patients for therapy. Across three highly pathogenic coronaviruses studied, SARS-CoV-2, MERS-CoV, and SARS-CoV-1, we find that allergic airway inflammation induced by HDM remodeling alters viral pathogenesis and disease severity. Unexpectedly, our data demonstrate divergent effects and outcomes, in which allergic airway inflammation is protective against the severity of response to SARS-CoV-2 and MERS-CoV infection, yet detrimental to SARS-CoV-1 infected mice. However, in all cases, HDM pre-treatment reduced lung viral titers by at least 10-fold.

Consistent with epidemiological studies from the COVID-19 pandemic [18–20, 39], our SARS-CoV-2 infection studies demonstrate that HDM induced allergic airway inflammation confers protection from weight loss, viral replication, lung immunopathology, and lethality. These protective effects were consistent across highly divergent SARS-CoV-2 strains including the Beta (B.1.351) variant, Omicron (BA.5) variant, and a lethal mouse adapted strain. This indicates a robust protective mechanism independent of virus strain differences.

Our experiments show that HDM treatment induces canonical allergic features including goblet cell hyperplasia, Muc5AC/Muc5B overexpression, M2a macrophage polarization, eosinophil recruitment, and IL-13 upregulation. Previously published *in vitro* data indicates that IL-13 can reduce ACE2 expression while increasing TMPRSS2 expression [31, 40], and that IL-13 treatment reduces SARS-CoV-2 replication in cell culture [41–43]. Our *in vivo* findings do not support an effect on either ACE2 or DPP4 receptor levels being responsible for differences in pathogenesis, since we did not find expression changes with HDM treatment in the lungs of mice.

We hypothesize that a combination of mucus accumulation, eosinophilia, and T2 immune skewing in HDM-treated mice collectively results in the alterations in pathogenesis of coronaviruses. The increase in mucins at the epithelial cell surface and T2 associated remodeling and inflammation might restrict viral access to ACE2-expressing cells, which results in suppressed spread. However, this model does not account for the observed increase in SARS-CoV- 1 pathogenesis and rapid destruction of epithelial cells [44]

Why is the outcome to SARS-CoV-1 infection different in HDM-treated mice than the other Betacoronaviruses we tested? First, there could be differences in intrinsic virulence and tropism, as SARS-CoV-1 exhibits greater lower airway replication and induces more potent innate immune activation, which may prime allergic lungs for uncontrolled inflammation [13, 45]. Second, there are distinct viral antagonism of host immune pathways among coronaviruses; SARS-CoV- 1 encodes strong interferon antagonists that could combine with T2 skewed immunity to create a hyperinflammatory, poorly regulated environment for host airways [46]. Third, there could be temporal differences between allergic inflammation and the more rapid SARS-CoV-1 replication kinetics we have previously observed in mice [29], such that T2 mediated remodeling and innate cell recruitment becomes detrimental rather than protective. Fourth, there may be enhanced susceptibility to immunopathology since SARS-CoV-1 infected HDM treated mice had higher chemokine levels associated with neutrophil, monocyte, and eosinophil recruitment, which are cell populations known to cause immune mediated lung injury in other contexts [44].

During the SARS-CoV-1 outbreak, few children were infected and pediatric cases were typically less severe [13, 14], mirroring observations from COVID-19. However, asthma prevalence among SARS-CoV-1 cases was not meaningfully evaluated, and experimental models were not examined. Our findings provide evidence that allergic airway inflammation might worsen SARS-CoV-1 pathogenesis *in vivo*, an observation potentially masked in human outbreaks due to limited case numbers in asthma populations and the rapid cessation of disease transmission. Conversely, our SARS-CoV-2 and MERS-CoV results are consistent with the epidemiological observation that allergic asthma does not worsen, and may even reduce, disease severity in these infections [18–21, 23, 47]

These findings suggest that allergic airway inflammation cannot be treated as a uniform risk factor across emerging coronaviruses. The data raises the possibility that host allergic airway status could influence transmission dynamics or viral evolution through the shifting of upper and lower airway replication niches. Further studies should dissect the factors distinguishing SARS-CoV-1, SARS-CoV-2, and MERS-CoV pathogenesis in allergic lungs, evaluate how different asthma endotypes (allergic, non-allergic, eosinophilic, neutrophilic) alter infection outcomes, and determine how airway remodeling impacts the tropism and immune evasion of newly emergent coronaviruses.

In SARS-CoV-2 and MERS-CoV, the antiviral effect, likely mediated by early viral control via eosinophils and mucins, modulates disease towards a milder outcome as mirrored by clinical and epidemiological data in humans. This same HDM mediated phenotype in SARS-CoV-1 changes the outcome from survival to lethality, likely mediated by immunopathology and airway damage induced by an extreme inflammatory response early in the course of infection. Future studies are warranted to characterize these mechanisms in further detail.

Importantly, the protective phenotype observed following HDM treatment is reproducible across multiple SARS-CoV-2 strains and distinct mouse genetic backgrounds, supporting the robustness of this response.

This study has several limitations. Although we demonstrate that HDM exposure differentially alters disease outcomes following infection with multiple highly pathogenic coronaviruses, the precise mechanisms underlying these virus-specific effects were not defined. Additional studies are needed to determine the relative contributions of altered epithelial responses, innate immune cell populations, and adaptive immunity to the observed phenotypes. Furthermore, these findings were generated in murine models of allergic airway inflammation and will require validation in human systems to determine their translational relevance.

Collectively, our findings reveal that allergic airway inflammation exerts virus-specific and divergent effects on disease severity of highly pathogenic coronaviruses, protective in both SARS-CoV-2 and MERS-CoV infection, yet pathogenic in SARS-CoV-1. These data enhance our understanding of host-virus interactions in chronic airway diseases and provide context for predicting risk and designing interventions for future coronavirus outbreaks.

## Materials and Methods

### *In vivo* mouse infections

All infections were performed in an animal biosafety level 3 facility at the University of Maryland, Baltimore, MD or Washington University School of Medicine in St. Louis, using appropriate practices, including a HEPA-filtered bCON caging system, HEPA-filtered powered air-purifying respirators (PAPRs), and Tyvek suiting. All animals were grown to 8+weeks of age prior to use in experiments. The animals were anesthetized using a mixture of xylazine (0.38 mg/mouse) and ketamine (1.3 mg/mouse) in a 50-μL total volume by intraperitoneal injection. The mice were inoculated intranasally with virus, after which all animals were monitored daily for weight loss and lung tissue was harvested for further analysis. All animals were housed and used in accordance with the University of Maryland, Baltimore, Institutional Animal Care and Use Committee guidelines.

### Four and six dose house dust mite model

House Dust Mite (HDM) Extract (Greer Labs, Lot #439881, endotoxin concentration: 523.628 EU/50 μg HDM dose) was resuspended in endotoxin-free PBS at 1 mg/mL. HDM-treated groups received 50 μg/treatment intranasally, while vehicle control groups received endotoxin-free PBS (Millipore-Sigma). Mice were treated on day zero and every 5^th^ day after for a total of 4 treatments as described previously [24].

Cohorts of 20-week-old male C57BL/6J mice were obtained from The Jackson Laboratory (000664). 25 µg of House Dust Mite extract (Greer labs, Lot #442871: 248.293 EU/25µg HDM dose) were resuspended in 50 mL of saline and given to mice via intranasal administration while control groups received saline only. Mice were treated with 3 doses of HDM extract per week for 2 weeks, followed by resting for 2 weeks, prior to virus inoculation.

### Mouse Infections with SARS-CoV-2 B.1.351 and BA.5 strains

BALB/c were obtained from The Jackson Laboratory (Bar Harbor, ME) and acclimated to specific pathogen-free conditions at the University of Maryland Baltimore. All animal experiments were age (8-10 weeks) and sex matched. The B.1.351 and BA.5 variants infection models of SARS-CoV-2 have been described previously [29, 30]. The viruses were passaged in Vero-TMPRSS2 cells and subjected to next-generation sequencing to confirm the stability of amino acid substitutions. Mice were euthanized on days 2 and 4 following infection with lung tissue collected for analysis or monitored for weight change and survival out to 4 days.

### Mouse infection with SARS-CoV-2 MA10

The MA10 stain of SARS-CoV-2 has been described [48]. The virus was passaged in Vero-TMPRSS2 cells and subjected to next-generation sequencing to confirm the stability of amino acid substitutions. Mice were inoculated with 3 x 10^5^ focus-forming units (FFU) of MA10. Weight and mortality events were recorded daily for 14 days. All MA10 experiments were performed in approved biosafety level 3 (BSL-3) facilities at Washington University School of Medicine. Animal studies were carried out in accordance with the recommendations in the Guide for the Care and Use of Laboratory Animals of the National Institutes of Health. The protocols were approved by the Institutional Animal Care and Use Committee at the Washington University School of Medicine (assurance number A3381–01). Virus inoculations were performed under anesthesia that was induced and maintained with ketamine hydrochloride and xylazine, and all efforts were made to minimize animal suffering. Sample size for animal experiments was determined using criteria set by the institutional Animal Care and Use Committee.

### Mouse infection with MERS-CoV

Mouse passaged MERS-CoV [37] was passaged in Vero-TMPRSS2 cells and subjected to next generation sequencing to confirm the stability of amino acid substitutions. Mice were inoculated with 5 x 10^3^ PFU of MERS-MA. Weight and mortality events were recorded daily for 10 days.

### Mouse infection with SARS-CoV-1

Mouse adapted SARS-CoV-1 MA-15 [29] was passaged in VeroE6 cells and subjected to next generation sequencing to confirm the stability of amino acid substitutions. Mice were inoculated with 1×10^2^-1×10^5^ PFU of MA-15. Weight and mortality events were recorded daily for 10 days.

### Serum HDM IgE detection

Mouse serum was collected prior to virus infection. Serum HDM-specific IgE was assessed using a commercial ELISA kit (Chondrex, 3037) and performed according to manufacturer’s instructions.

### Histology

Mouse lungs were fixed 10% formalin in PBS for at least 48 hours. These lungs were then processed at the UMB Histology core facility for paraffin embedding, sectioned into 5 mm sections, and stained with hematoxylin and eosin staining.

### Immunohistochemistry

Paraffin-embedded and sectioned mouse lungs were used for rat anti-mouse F4/80 immunostaining (Thermo Fisher, #14-4801-82). Paraffin-embedded and sectioned lungs were used for MUC5AC and MUC5B immunohistochemistry staining. For MUC5AC staining, a primary Muc5AC monoclonal antibody (Invitrogen, INV-MA5-12178) at a 1:100 dilution, and a secondary M.O.M Biotinylated Anti-Mouse IgG reagent (Vector Laboratories, RND-BMK-2022- NB) with Basic Immunodetection Kit. For MUC5B staining, a primary Polyclonal antibody (Invitrogen, PA5-142969) at a 1:1000 dilution, and a Secondary antibody, Vectastain Elite ABC anti-Goat IgG HRP Immunodetection kit (Novus Biologicals, PK-6105-NB). Mucin and YM1 staining were quantified as a stained percentage of the total field performed three times per lung and averaged using ImageJ software.

### RNA sequencing

Transcriptomic libraries were prepared and sequenced by the Yale Center for Genome Analysis (West Haven, CT, USA) using a modified version of a previous analysis workflow [49]. Raw data are available at the BioProject accession number PRJNA1389634. Read preprocessing was performed using cutadapt v3.4, followed by alignment to the to the murine genome (assembly GRCm38) using STAR v2.7.8a[50]. DESeq2 v4.1.0[51] was used to perform gene expression analysis in R (Rstudio; Boston, MA, USA). The resultant differential expression matrices were used to perform pathway analysis using Ingenuity Pathway Analysis (QIAGEN, Hilden, Germany) using an alpha value of padj < 0.01 to classify a gene as significantly differentially expressed.

### Virus titering

Vero-E6/TMPRSS2 cells were cultured in DMEM (Dulbecco’s Modified Eagle Medium) (Quality Biological), supplemented with 10% FBS (Sigma), 1% (vol/vol) penicillin/streptomycin (Gemini Bio-products), and 1% (vol/vol) L- glutamine (Gibco). Cells were grown and maintained in an incubator at 37°C and 5% CO2. Viral lung titers were quantified by homogenizing lung tissues in PBS (Quality Biological) with 1.0 mm glass beads (Sigma) in a Bead raptor (Omni International). Vero-E6 cells were plated in 12-well plates with 2.0 × 10^5^ cells per well. Plaque assays were performed by adding 25 μL of lung homogenate after centrifugation to 225 μL of DMEM with 10-fold dilutions across a six-point dilution curve with 200 μL of DMEM diluent added to each of the wells. After a 1-h dilution with plate rocking every 15 min, a 2 mL agar overlay containing DMEM was added to each of the wells. Plates were incubated for 2 days for all viruses at 37°C and 5% CO2 and then fixed with 10% formalin, stained with crystal violet, washed with tap water, and plaques were counted.

### Cytokine and chemokine quantification

Lung tissue was homogenized, and a Bio-Plex Pro Mouse Chemokine Panel multiplex bead array assay (BIORAD) was used to quantify an array of cytokines and chemokines as listed in the manufacturer protocol. An additional fixation step (10% formalin (Sigma) at 4°C x 12 h) was added at the end of the protocol following the last wash. The plate was then washed three times and analyzed on a Luminex MagPix and xPONENT Software version 4.3. Using a standard curve, concentrations of each analyte were calculated for all samples.

### Gene expression by qPCR

Mouse lung tissue was homogenized in 1 mL of TRIzol (Ambion) using 1.4 mm glass beads (Omni International) and a Beadruptor (Omni International). RNA was extracted using the DIRECT-zol mini-RNA Extraction kit (Zymo Research) per the manufacturer’s instructions. mRNA extraction was followed by cDNA reverse transcription and quantitative PCR. Total purified RNA was reverse transcribed to cDNA, and real-time qRT-PCR was performed for *Muc4, Muc1, Muc5ac, Muc5b, Ace2, IL-4, IL-13, YM1, Arg1,* and *RelmA. Gapdh* was used as a housekeeping gene for normalization. Results were confirmed as fold changed (2^-ΔΔCT) in mRNA expression with relative expression normalized to uninfected vehicle groups.

### Neutrophil depletion

Neutrophils were depleted with 500 µg of anti-Ly6G (clone 1A8) or isotype antibody (IgG2a isotype) (Bio-X-Cell, Lebanon, NH), *via* intraperitoneal injection 1 day prior to infection with virus and every 2 days afterward. Mice were infected with either 50 µL sterile PBS or 10^3^ PFU of SARS-CoV-1 MA-15. Mice were monitored daily for weight loss and signs of morbidity. On day 7, animals were euthanized, and lung tissue was collected for further analysis.

### Flow Cytometry

Single cells were isolated from the lungs using the Lung Dissociation Kit, mouse and gentleMACS Dissociator (Miltenyi Biotec). Lungs were placed in C tubes (Miltenyi Biotec) containing 2.2 mL Buffer S, 15 µL of Enzyme A, and 100 µL of Enzyme D. Lungs were digested on a gentleMACS (Milteni) for 30 min at 37°C. Cell suspensions were filtered on a 70 µm filter (BD), and cells were pelleted by centrifugation (300 x g for 10 min). Red blood cells were lysed in ammonium-chloride-potassium lysis buffer (Quality Biological, Inc.) and subsequently washed with PBS containing 3% FBS. Approximately, 10^6^ cells were plated and washed twice with PBS containing. Cells were stained for viability using the Live/Dead Fixable NIR Dead Cell Stain Kit (Molecular Probes). Cells were washed with PBS containing 3% FBS. Cells were stained with antibody cocktails made in BD Horizon Brilliant Stain Buffer (BD): CD45 Alexa Fluor 700 (1:200; BioLegend, clone 30-F11), CD11b Brilliant Violet 650 (1:167; BioLegend, clone M1/70), Ly6G Brilliant Violet 785 (1:100; BioLegend, clone 1A8), Ly6C Brilliant Violet 510 (1:20; BioLegend, clone HK1.4), CD3 BUV563 (1:20; BD, clone 145- 2C11), CD4 FITC (1:200; BioLegend, clone GK1.5), CD8a APC (1:200; BioLegend, clone 53-6.7), CD19 PE (1:50; BioLegend, clone 6D5), NK1.1 Pacific Blue (1:200; BioLegend, clone PK136), Siglec-F Brilliant Violet 605 (1:20; BioLegend, clone X54-5/7.1), CD103 BUV661 (1:20; BD, clone 2E7), Siglec-H BUV563 (1:20; BD, clone 440c), and Siglec-F Brilliant Violet 711 (1:100; BD, clone E50-2440). Cells were sorted on Cytek Aurora-5 machine and analyzed using FlowJo V10 software.

### Statistical analysis

Statistical analyses were performed with GraphPad Prism software (GraphPad Software, San Diego, CA). Data were analyzed using unpaired *t-test*, one-way or two-way analysis of variance (ANOVA) followed by Tukey, Dunnett, or Sidak post hoc comparison test as indicated. All statistical analyses were two-sided and a *P* < 0.05 was considered statistically significant. For the survival study, the Kaplan-Meier survival curve was used, and a log-rank (Mantel-Cox) test was performed to determine significance. Statistical methods were not used to predetermine the sample size. Blinding and randomization were not used.

## Data availability

All data supporting the findings of this study are included in the manuscript and supplementary material. Raw RNA sequencing data have been deposited in the NCBI BioProject database under accession number PRJNA1389634. Raw data is available at 10.5281/zenodo.22130998 or through contact with Matthew Frieman.

## Acknowledgments

We thank Lisa Hester of the University of Maryland School of Medicine Cytokine Core Laboratory for assistance in optimizing and running Luminex plates for Bioplex assays, and Perry Comegys and Norma Colocho for assistance with embedding, sectioning, and staining of lung slides. Multiple figures contained images created with Biorender.com.

